# Circadian-Modulated Thresholds as a Mechanistic Basis for Sleep-Wake Transitions, Recovery, and Sleepiness

**DOI:** 10.64898/2026.02.04.703688

**Authors:** Chenggui Yao, Junjie Jiang, Changgui Gu, Jianwei Shuai, Dongping Yang

## Abstract

Human sleep-wake cycles arise from the interplay between homeostatic sleep pressure and circadian rhythms, yet the underlying mechanistic basis by which these drives jointly govern state transitions, recovery from sleep loss, and subjective sleepiness remains unclear. Here, we use the extended Phillips-Robinson model incorporating circadian excitation of orexin and locus coeruleus populations, yielding analytically tractable, circadian-modulated thresholds for sleep onset and awakening, and delineating the roles of circadian and homeostatic drives. We show that homeostatic feedback alone generates intrinsic sleep-wake oscillations via a saddle-node on invariant circle bifurcation, while circadian drive reshapes the stability landscape to account for immediate sleep onset and partial first-night recovery after prolonged deprivation, and enables analytic predictions of sleep timing and duration. We further define sleepiness as the distance between the homeostatic state and the active circadian sleep threshold, which robustly predicts subjective sleepiness across various deprivation, restriction, extension, and recovery protocols. Together, these results establish circadian-modulated thresholds as a unifying dynamical principle linking sleep-wake transitions, recovery dynamics, and sleepiness, with implications for circadian misalignment, shift work, and individualized sleep interventions.

**Author summary:** Sleep and wakefulness arise from the interaction between circadian timing and the gradual accumulation and dissipation of sleep pressure. Although existing computational models have advanced understanding of these processes, most rely on fixed or heuristic rules for switching between sleep and wake states, limiting their ability to explain behavior under sleep deprivation, extension, or restriction. Here, we present a mechanistic description of state switching in which circadian and homeostatic influences act as separable but interacting control dimensions, providing a framework for how circadian modulation shapes state stability, recovery sleep, and subjective sleepiness, without invoking separate mechanisms for normal and perturbed conditions. By recasting sleep regulation in terms of state-dependent dynamical boundaries, our results offer a unified perspective on sleep timing, duration, recovery, and vulnerability to fatigue. This work provides a more physiologically grounded foundation for sleep-wake modeling and a principled basis for understanding circadian misalignment, shift work, jet lag, and other real-world challenges to sleep health.

## Introduction

Sleep and wakefulness are fundamental behavioral states governed by the interaction of two core physiological processes [1]: a circadian rhythm with an approximately 24-hour period and a homeostatic sleep pressure that accumulates during wakefulness and dissipates during sleep. Borbély’s seminal two-process model formalized this interaction by proposing that sleep–wake timing emerges from circadian-dependent thresholds for sleep onset and awakening [2, 3], providing a unifying account of a wide range of behavioral and physiological observations. However, these thresholds were originally introduced in an *ad hoc* manner, without explicit mechanistic grounding [4, 5]. Subsequent experimental and computational studies have demonstrated that circadian signals influence sleep–wake regulation through multiple physiological pathways, including direct modulation of wake- and sleep-promoting neural populations and their mutual inhibition [6–9], thereby providing a physiological basis for circadian-dependent sleep and wake thresholds and motivating their explicit incorporation into dynamical models of arousal-state regulation.

The suprachiasmatic nucleus (SCN), the brain’s central circadian pacemaker, exertsm time-of-day–dependent influences on sleep- and arousal-regulating circuits, including direct and indirect projections to the ventrolateral preoptic area (VLPO) and other hypothalamic and brainstem nuclei [10, 11]. In parallel, the SCN coordinates rhythmic endocrine outputs—most prominently melatonin and cortisol—that modulate neuronal excitability and systematically bias the propensity for sleep or wakefulness across the circadian cycle [12–14]. Circadian control of thermoregulatory processes, particularly the rhythm of core body temperature, further contributes to the timing of sleep onset and the stability of sleep [12, 15]. Crucially, circadian phase gates the system’s sensitivity to homeostatic sleep pressure (e.g., adenosine), such that identical levels of accumulated sleep drive can produce qualitatively different behavioral outcomes depending on circadian time [13]. Together, these convergent pathways establish a physiological basis for modeling sleep and wake thresholds as dynamically modulated quantities rather than arbitrarily imposed constants. The Phillips–Robinson (PR) model formalizes this principle by introducing two biologically interpretable, circadian-dependent thresholds that govern transitions between sleep and wake states [7, 9].

These thresholds have proven essential for accurately capturing sleep-wake dynamics and have enabled a range of practical and translational applications. For example, Shochat *et al*. used model-derived sleep and wake thresholds to examine the relationship between sleep timing and subjective sleepiness, quantified by the Karolinska Sleepiness Scale (KSS) [16], showing that elevated evening sleepiness was associated with earlier bedtimes and longer subsequent sleep duration [17]. Building on this framework, mathematical models incorporating circadian-dependent thresholds have been used to design personalized light-based interventions that optimize sleep timing, particularly in aging populations [18]. The Homeostatic–Circadian–Light model further demonstrated that explicitly modeling both sleep and wake thresholds substantially improves predictions of sleep timing and duration across varying environmental light conditions [19]. More recently, Hong *et al*. introduced the concept of circadian sleep sufficiency (CSS), a threshold-based metric integrating circadian modulation of sleep propensity, to quantify and mitigate excessive daytime sleepiness in shift workers [20, 21]. Collectively, these advances highlight the central role of biologically interpretable thresholds in sleep-wake models and underscore their translational value for predicting sleepiness, optimizing interventions, and improving sleep health across diverse populations.

Despite substantial progress, a key limitation of the PR model is that circadian modulation of the sleep–wake switch is conveyed solely via input to the VLPO [10, 22], neglecting well-established excitatory circadian projections to locus coeruleus (LC) and orexin/hypocretin (Orx) neurons [23, 24]. Converging experimental evidence shows that SCN projections to Orx and LC are critical for stabilizing sleep–wake transitions [24, 25], and that robust circadian regulation persists even after VLPO lesions [26–28], indicating that SCN-mediated inhibition of VLPO and excitation of Orx/LC act in parallel to shape sleep–wake dynamics. Existing theoretical models have emphasized circadian excitation of wake-active populations [29–31], successfully accounting for diurnal, nocturnal, and bimodal sleep–wake patterns [29] and for the stabilizing role of Orx in sleep–wake transitions [30]. However, these circadian wake-promoting drives are typically explored under idealized or phenomenological conditions and remain largely disconnected from real-world measures of alertness or subjective experience [29–31]. In contrast, subjective sleepiness—most commonly quantified using the KSS–is central to translational applications such as fatigue risk management, clinical assessment, and intervention design, yet existing circadian–homeostatic models lack a direct quantitative connection between circadian-modulated wake drive, transition thresholds, and KSS. This gap motivates the development of a unified dynamical framework that explicitly integrates parallel circadian pathways, physiologically grounded thresholds, and subjective sleepiness within a single mechanistic model.

To address this gap, we extend the PR framework by explicitly incorporating circadian excitatory drive to Orx/LC populations, thereby redefining the dynamical architecture of sleep–wake switching. This yields analytically tractable, circadian-modulated sleep and wake thresholds that specify when state transitions occur and disentangle circadian and homeostatic control at the dynamical level. We show that homeostatic pressure alone generates intrinsic oscillations via a saddle-node on invariant circle (SNIC) bifurcation, while circadian drive reshapes the stability landscape by deforming these thresholds. Within the same geometric framework, the model reproduces immediate sleep onset after extreme deprivation and partial first-night recovery after chronic deprivation, jointly constraining the strength of circadian excitation of wake-promoting populations. Finally, the explicit thresholds enable a mechanistic definition of sleepiness as the distance to the circadian sleep boundary, which accurately predicts subjective sleepiness across deprivation, restriction, extension, and recovery protocols. Together, these results establish circadian-modulated thresholds as a unifying dynamical principle linking state transitions, recovery dynamics, and sleepiness.

### Flip-Flop Sleep-wake Model with Circadian-modulated Wake- and Sleep-Drives

We extend the physiologically grounded PR model by incorporating excitatory circadian drive to wake-promoting neurons (Fig. 1). The model centers on a bistable flip-flop architecture comprising inhibiting mutually sleep-active and wake-active neural populations, whose dynamics are jointly regulated by circadian and homeostatic processes [32–34]. Circadian timing (Process C) originates in the SCN, which entrains internal rhythms to the light-dark cycle and modulates sleep-wake state through downstream hypothalamic pathways. In addition to the well-established inhibitory influence on the VLPO via the dorsomedial hypothalamus (DMH) [10], anatomical and functional evidence indicates that the SCN also provides the excitatory input to wake-active Orx/LC neurons [24, 25], as shown in Fig. 1a. This excitatory circadian projection, omitted in the original PR model, is explicitly incorporated here and provides a parallel pathway for circadian control of arousal. For analytical simplicity, the direct inhibition from VLPO to Orx/LC neurons as explicitly modeled in [30] is subsumed within the effective flip-flop architecture. Homeostatic sleep drive (Process S) captures the accumulation and dissipation of sleep pressure, rising during wakefulness and declining during sleep [35, 36].

**Fig 1.**
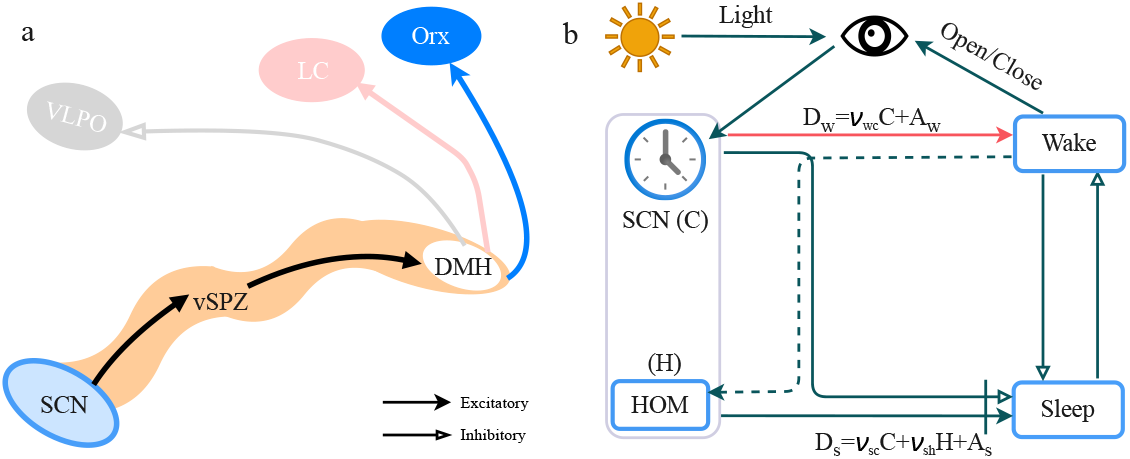
Circadian control of the sleep–wake switch and model architecture. (a) Anatomical pathways mediating circadian regulation of sleep and wakefulness. Output of the SCN is relayed via the ventral and dorsal subparaventricular zones (vSPZ and dSPZ) to the DMH, which in turn modulates sleep-promoting neurons in VLPO and wake-promoting Orx/LC neurons. (b) Schematic of the modified sleep-wake model based on the PR model [32]. In addition to circadian inhibition of the VLPO, the model incorporates an explicit excitatory circadian drive to wake-promoting Orx/LC neurons (red arrow), reshaping the circadian-dependent modulation of sleep and wake thresholds.

It is mediated by sleep-promoting neuromodulators, notably adenosine and prostaglandin D_2_ (PGD_2_), which act on the VLPO and related sleep-active circuits to bias the sleep-wake switch toward sleep [37–40]. In the model, sleep-wake transitions emerge from the interaction between Process S and circadian modulation (Process C) through circadian-dependent thresholds: increasing homeostatic pressure triggers sleep when sleep-active populations dominate (sleep branch in the saddle-node bifurcation), whereas decreasing pressure permits wakefulness when wake-active populations prevail (wake branch in the saddle-node bifurcation) [9]. Incorporating circadian excitation of Orx/LC neurons refines these dynamic, state-dependent thresholds, providing a unified framework for analyzing sleep-wake transitions, sleep deprivation, and sleepiness.

Figure 1(b) summarizes the model architecture. The dynamics of the wake-active population (*w*), sleep-active population (*s*), and homeostatic pressure (*H*) are governed by:

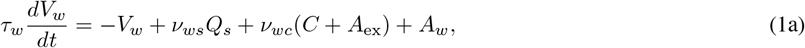

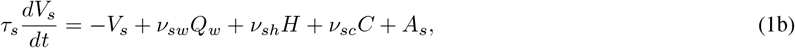

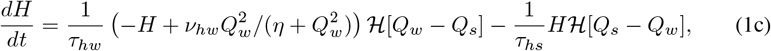

where *V*_*a*_ (mV) (*a* = *w, s*) denotes the mean cell-body potential of each population, and the firing rate *Q*_*a*_ is given by a sigmoid function:

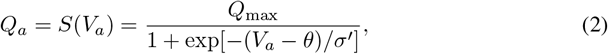

with *S*(·) being the sigmoidal function, *Q*_max_ the maximum firing rate, *θ* the mean firing threshold relative to resting, and 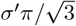 the deviation [32]. The time constants *τ*_*a*_ (*a* = *w, s*) represent the characteristic neuromodulatory decay times. Mutual inhibition between populations is modeled by negative coupling coefficients *v*_*ws*_ and *v*_*sw*_. Constant inputs *A*_*a*_ (*a* = *w*, ex, *s*) represent time averaged inputs to each population from external sources and could include any constant offsets of the circadian drive:

*A*_*w*_ models cholinergic input to wake-active neurons, *A*_ex_ captures constant part of excitation from Orx and LC populations, and *A*_*s*_ is set to zero.

For analytical tractability, the circadian drive is modeled as a sinusoidal signal, *C*(*t*) = sin(*ω*_*c*_*t*) with 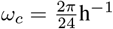, consistent with prior theoretical work [41, 42]. Its amplitude is normalized, with effective strength absorbed into the coupling coefficients. The homeostatic drive (*H*), measured in percent slow-wave activity (SWA) power [37, 38], increases (with *v*_*hw*_ > 0) during wake and decreases during sleep. Accumulation is governed by a nonlinear, saturating function of wake firing rate *Q*_*w*_, reflecting sleep-substance buildup (e.g., adenosine), while dissipation occurs exponentially during sleep. State dependence is enforced by the Heaviside function ℋ [.], with transitions determined by the relative dominance of wake- and sleep-active firing [30]. The parameter *v*_*sh*_ > 0 quantifies the excitatory influence of the homeostatic process on sleep-active neurons, consistent with experimental and modeling studies showing that elevated sleep pressure directly facilitates VLPO activation [29, 32]. The time constants *τ*_*hw*_ and *τ*_*hs*_ determines the rates of homeostatic pressure accumulation during wake and dissipation during sleep, respectively, and are set to experimentally constrained values characteristic of adult human sleep behavior [43]. All parameters with nominal values are summarized in Tab. 1.

**Table 1.** Nominal parameter values.

| Parameter | Value | Unit | Description |
| --- | --- | --- | --- |
| $\nu_{ws}$ | -2.0 | mV s | Input strength to wake neurons from sleep neurons |
| $\nu_{sw}$ | -2.0 | mV s | Input strength to sleep neurons from wake neurons |
| $\nu_{wc}$ | $0 \sim 3$ | mV | Excitatory input strength to wake neurons from SCN |
| $\nu_{sc}$ | -0.3 | mV | Inhibitory input strength to sleep neurons from SCN |
| $\nu_{sh}$ | $0 \sim 3$ | mV nM<sup>-1</sup> | homeostatic input strength to sleep neurons |
| $\tau_w$ | 10 | s | Decay time constant of wake neuromodulator |
| $\tau_s$ | 10 | s | Decay time constant of sleep neuromodulator |
| $A_w$ | 1.4 | mV | Constant input from the acetylcholine groups |
| $A_{ex}$ | 1.0 | | Constant input from the Orexin and LC groups |
| $A_s$ | 0 | mV | Constant offset of the circadian drive |
| $\tau_{hw}$ | 45 | h | Increase time constant of homeostatic sleep pressure |
| $\tau_{hs}$ | 4.8 | h | Decay time constant of homeostatic sleep pressure |
| $\nu_{hw}$ | 20 | nM | Upper bound for homeostatic sleep pressure |
| $\eta$ | 2.3 | s<sup>-2</sup> | Control parameter of homeostatic production |
| $Q_{\max}$ | 100 | s<sup>-1</sup> | Maximum possible firing rate of neurons |
| $\theta$ | 10 | mV | Mean firing threshold relative to resting |
| $\sigma'$ | 3 | mV | Standard deviation for mean firing threshold |

Compared to the original PR model, the key modification is the introduction of an excitatory circadian input to the wake-promoting population (the term *v*_*wc*_(*C* + *A*_*ex*_)), representing SCN-driven excitation of Orx and LC neurons. Together with circadian inhibition of VLPO (*v*_*sc*_), this creates parallel excitatory and inhibitory circadian pathways that dynamically shape sleep and wake thresholds. This formulation enables systematic analysis of how circadian drive and homeostatic pressure jointly regulate state transitions, sleep deprivation responses, and sleepiness. Parameter values in Tab. 1 are chosen to remain consistent with prior flip-flop models [32, 44] and experimental observations under circadian misalignment and sleep loss [30, 45].

## Results

### Dynamical Mechanism of Sleep-Wake Cycles

We first examined how the interaction between homeostatic sleep pressure and circadian excitation shapes intrinsic sleep-wake dynamics in the modified model. Figure 2a shows the activity of wake- and sleep-promoting populations (*Q*_*w*_, *Q*_*s*_) with various homeostatic gains *v*_*sh*_ and one fixed circadian excitation to wake-active neurons (*v*_*wc*_). Four qualitatively distinct regimes emerge. For low *v*_*sh*_, sleep pressure accumulates insufficiently, leading to prolonged wakefulness and failure of sleep initiation. At intermediate *v*_*sh*_, short and unstable sleep episodes occur, reflecting weak homeostatic drive that cannot sustain consolidated sleep. At nominal values, the system exhibits a stable biphasic rhythm with approximately 16 h of wakefulness followed by 8 h of sleep, consistent with healthy adult sleep patterns. For large *v*_*sh*_, excessive sleep pressure fragments wakefulness and produces polyphasic sleep-wake dynamics. Thus, modulation of homeostatic gain alone, in the presence of circadian excitation, reproduces a wide spectrum of sleep phenotypes spanning normal and pathological regimes, highlighting the central role of homeostatic–circadian interaction in shaping sleep-wake stability.

**Fig 2.**
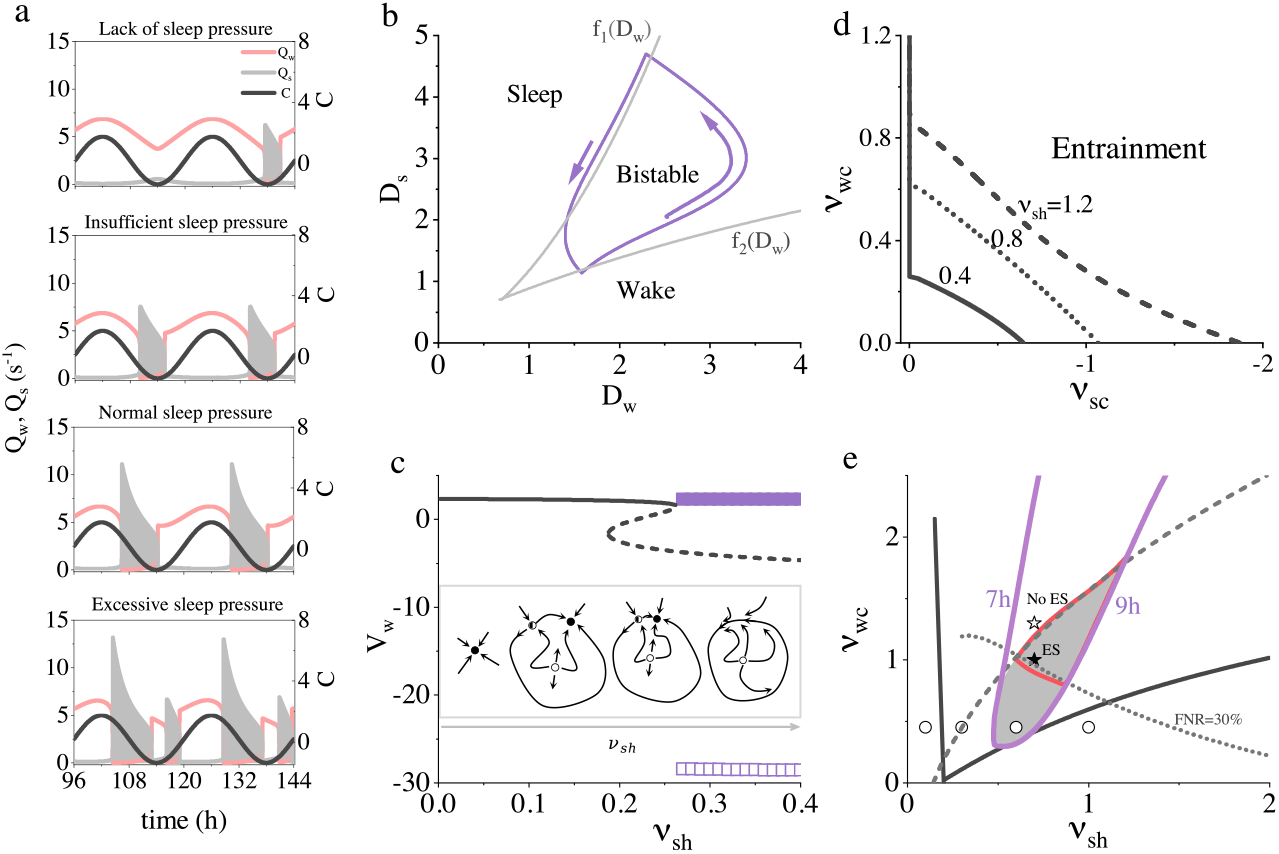
Dynamical mechanism of sleep-wake cycles. (a) Time series of wake (*Q*_*w*_) and sleep (*Q*_*s*_) activity for four values of homeostatic gain *v*_*sh*_ = 0.1, 0.3, 0.6, 1.0 with circadian input fixed at *v*_*wc*_ = 0.45 (denoted as open circles in (e)). Increasing *v*_*sh*_ leads to transitions from prolonged wakefulness to polyphasic sleep patterns. (b) Dynamical phase diagram on (*D*_*w*_, *D*_*s*_) plane: wake, sleep, and bistable regions. Nominal trajectory forms a sleep-wake cycle, crossing sleep-onset and awakening boundaries: *D*_*s*_ = *f*_1_(*D*_*w*_) and *D*_*s*_ = *f*_2_(*D*_*w*_), which are analytically derived as Eq. 9. (c) Bifurcation diagram showing steady-state and oscillatory values of *V*_*w*_ versus *v*_*sh*_, in the absence of the circadian drive. An SNIC bifurcation gives rise to a limit cycle, marking the onset of rhythmic sleep-wake behavior (maxima: filled; minima: open squares), where the stable wake state (solid line) transitions become unstable (dashed line). Inset: phase space schematics showing the evolution of fixed points (stable: filled; saddle with one positive and two negative eigenvalues: half-filled; saddle with two positive and one negative eigenvalues: open circles; the manifold connecting different states: dashed line). (d) Entrainment regions in the (*v*_*wc*_, *v*_*sc*_) plane at various *v*_*sh*_. (e) Phase diagram in the (*v*_*sh*_, *v*_*wc*_) plane. Black line: entrainment boundary; Purple-outlined region: normal sleep duration (7∼ 9 h); Gray region: Enter Sleep (ES) regime, characterized by immediate sleep onset following acute sleep deprivation, whose upbound is predicted by theoretical analyses (dashed line, Eq. 16); Red-outlined region: First Night Recovery (FNR), in which partial recovery sleep occurs during the first post-deprivation night, accounting for approximately 10 ∼ 30% of the accumulated sleep loss, whose lower bound (FNR=30%) is predicted by theoretical analyses (dotted line, Eq. 21). Stars: two deprivation cases in Fig. 4c.

Although circadian rhythms are often interpreted through the lens of external entrainment, the intrinsic dynamical mechanism by which circadian and homeostatic processes jointly generate robust cyclic transitions remains incompletely understood. Previous studies [30, 31, 46] addressed this problem primarily through fast–slow manifold reductions 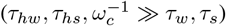, in which state switching is attributed to saddle-node bifurcations of the fast neural subsystem driven by the net synaptic inputs

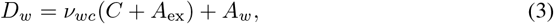

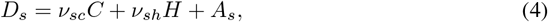

which defines a two-dimensional slow phase space for analyzing sleep-wake thresholds and transitions (Fig. 2b). While this reductionist framework successfully captures the existence of transitions, it fundamentally merges circadian (*C*) and homeostatic (*H*) influences into aggregated drive variables, thereby precluding a principled characterization of their distinct dynamical roles in shaping stability and switching.

To overcome this limitation, we explicitly decoupled the two drives by isolating the homeostatic component (setting *v*_*sc*_ = *v*_*wc*_ = 0) and performing a full-system bifurcation analysis without imposing timescale separation, following our previous work [47]. As *v*_*sh*_ increases, the system transitions from a stable fixed point corresponding to tonic wakefulness to sustained oscillations via a SNIC bifurcation (Fig. 2c). At the bifurcation, a stable node, saddle, and unstable node coalesce into a heteroclinic loop, from which a stable limit cycle emerges (Fig. 2c, inset). This SNIC mechanism provides a biologically grounded explanation for how increasing homeostatic pressure alone can generate rhythmic sleep-wake alternations, establishing a dynamical substrate upon which circadian modulation subsequently acts.

We next investigated entrainment to a 24-h light–dark cycle by varying the circadian drive gains to sleep- and wake-promoting populations, *v*_*sc*_ and *v*_*wc*_. Stable synchronization produces a robust 24-h rhythm over an extended region of the (*v*_*wc*_, *v*_*sc*_) plane (Fig. 2d). When *v*_*wc*_ = 0, entrainment requires substantially larger *v*_*sc*_, and the critical circadian strength increases with homeostatic pressure, consistent with the original PR model [7]. In contrast, increasing *v*_*wc*_ markedly expands the entrainment region and lowers the minimal circadian drive required for synchronization, enabling stable entrainment even for *v*_*sc*_ = 0. These results demonstrate that coordinated circadian modulation of both sleep- and wake-promoting circuits substantially enhances the robustness and flexibility of entrainment to environmental forcing.

Complementarily, when *v*_*sc*_ is held fixed, the system exhibits an Arnold-tongue–shaped entrainment region in the (*v*_*wc*_, *v*_*sh*_) plane (Fig. 2e), revealing a nontrivial interaction between circadian wake drive and homeostatic sleep pressure. Within this entrainment domain, additional physiological constraints further restrict the admissible parameter space, including normal sleep duration (7 ∼ 9 h), immediate sleep onset following acute deprivation (ES), and partial first-night recovery after chronic deprivation (FNR ∼10 ∼30%; Sec.). Together, these results show that successful circadian entrainment and physiologically realistic sleep-wake dynamics are jointly determined by the balance between circadian excitation of wake-promoting populations and homeostatic sleep pressure, and that explicitly incorporating both SCN pathways substantially enlarges the biologically plausible operating regime.

### Dynamical Interaction between Circadian and Homeostatic Drives

We now characterize how circadian and homeostatic processes jointly shape state-transition thresholds, and thereby determine sleep timing and duration. Unlike the above analyses that focus primarily on oscillation generation or entrainment boundaries, here we examine how circadian modulation reshapes homeostatic switching thresholds. Exploiting the strong timescale separation 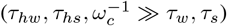, the wake- and sleep-promoting populations can be treated as fast variables responding quasi-statically to slowly varying drives (*D*_*w*_, *D*_*s*_), yielding the reduced two-dimensional fast subsystem

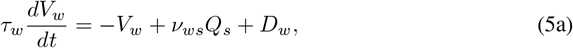

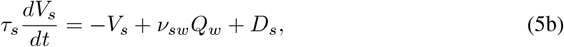

whose fixed points and bifurcations define the instantaneous stability of wake or sleep state. Theoretical analysis yields sleep-onset threshold *D*_*s*_ = *f*_1_(*D*_*w*_) and awakening one *D*_*s*_ = *f*_2_(*D*_*w*_) (Fig. 2b), and, by projection, explicit circadian-dependent thresholds for the homeostatic sleep pressure: *H*^+^ for sleep onset and *H*^*−*^ for awakening (see Sec. for details).

Importantly, when the mutual inhibition strengths satisfy *v*_*ws*_ = *v*_*sw*_, these two boundaries are exact inverses in the (*D*_*w*_, *D*_*s*_) plane, reflecting an intrinsic symmetry between sleep- and wake-active populations (Fig. 2b). In this regime, system trajectories drift slowly under circadian and homeostatic modulation and undergo rapid transitions upon crossing the fold curves. While classical fast–slow analyses describe similar fold-mediated transitions [30, 46], the present formulation makes two advances. First, circadian and homeostatic influences remain explicitly parameterized rather than being absorbed into undifferentiated control variables: distinct parameter combinations (*v*_*sh*_, *v*_*sc*_, *v*_*wc*_) generate slow trajectories in the (*D*_*w*_, *D*_*s*_) plane, while the analytically derived sleep and wake thresholds correspond precisely to the fold curves of the fast subsystem. Second, the transition boundaries are obtained in closed form, enabling direct quantitative predictions of how physiological parameters modulate sleep onset and awakening. In the spectial case *v*_*wc*_ = 0, these expressions reduce to the thresholds of the original PR model [20, 32]. For *v*_*wc*_ > 0, however, the thresholds acquire a nonlinear dependence on circadian amplitude, reflecting the role of the excitatory SCN input to Orx/LC neurons and providing a mechanistic account of circadian modulation of state transitions beyond VLPO-mediated inhibition alone. Thus, circadian input does not merely bias state durations, but actively reshapes the switching geometry by dynamically repositioning the homeostatic thresholds.

Figure 3a illustrates these dynamics by plotting the homeostatic sleep pressure *H* (black traces) together with the analytically derived, circadian-dependent thresholds *H*^+^ and *H*^*−*^ (Eq. 10). Sleep occurs when *H* exceeds the sleep-onset threshold *H*^+^ (orange solid line), whereas wakefulness resumes when *H* falls below the awakening threshold *H*^*−*^ (orange dash-dotted line). Increasing the homeostatic coupling strength *v*_*sh*_ amplifies the accumulation and impact of sleep pressure, prolonging sleep episodes by sustaining activation of sleep-promoting populations (Fig. 3b, left). By contrast, increasing the circadian excitation of wake-active neurons (*v*_*wc*_) elevates both thresholds, thereby delaying sleep onset, extending wake duration, and reducing total sleep time (Fig. 3b, right). Systematic parameter sweeps in the (*v*_*wc*_, *v*_*sh*_) plane delineate regimes consistent with physiologically normal sleep durations (7∼ 9 h; Fig. 3b and purple boundary in Fig. 2e), indicating that appropriate balance between circadian excitation and homeostatic drive is required for realistic sleep–wake organization. Sleep timing is shaped in a similarly structured manner (Fig. 3c): stronger homeostatic coupling advances transitions and biases the system toward longer sleep, whereas stronger circadian excitation of wake-active neurons shifts both sleep onset and offset to later circadian phases.

**Fig 3.**
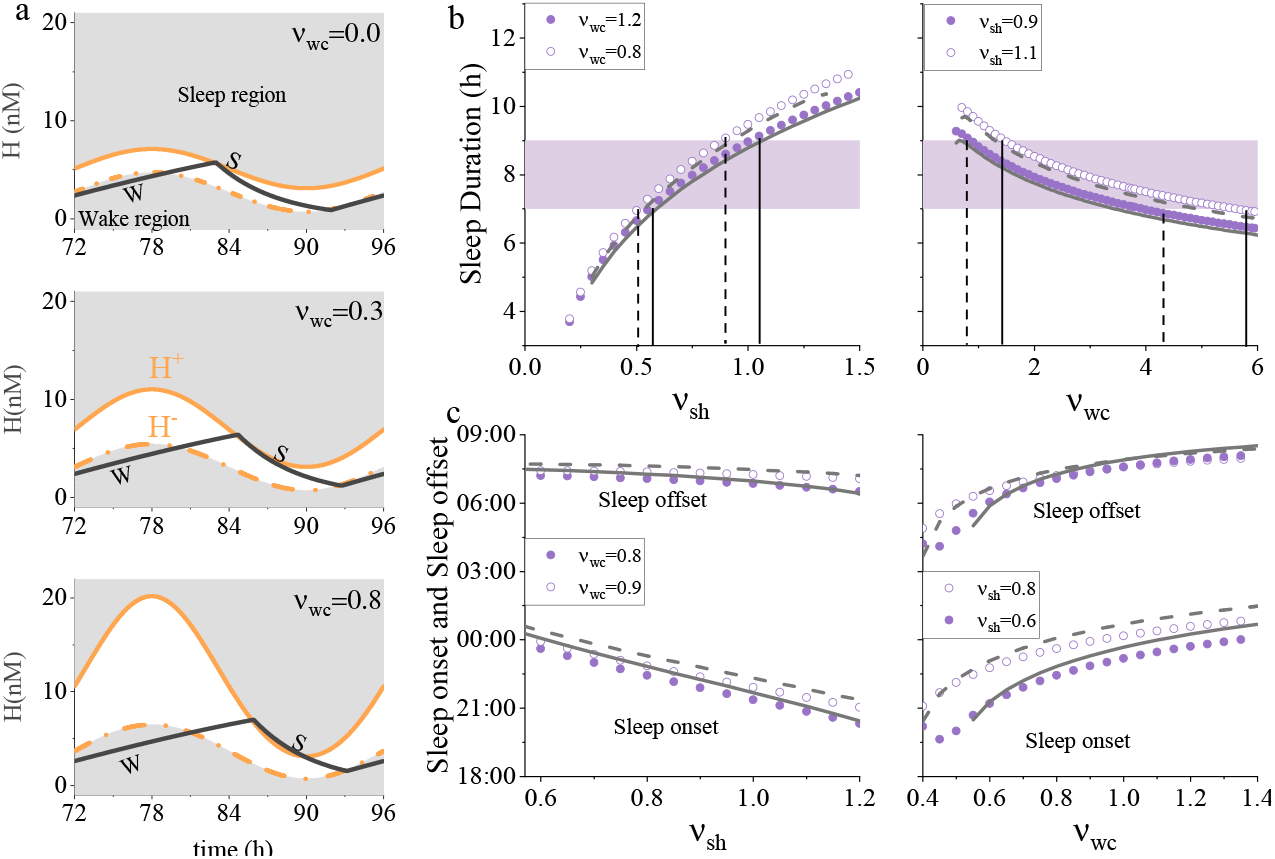
(a) Interaction between homeostatic sleep pressure (black line) and circadian sleep-wake boundaries (orange lines) at various gains *v*_*wc*_. solid line: sleep threshold (H^+^), dash-dotted line: wake threshold (H^*−*^). (b) Sleep duration *T*_*s*_ against *v*_*sh*_ (left) and *v*_*wc*_ (right). Simulation and analytical results (Eq. 11) are denoted by circles and lines, respectively. Open circles and dashed line: *v*_*wc*_ = 0.8 at left and *v*_*sh*_ = 1.1 at right; Filled circles and solid line: *v*_*wc*_ = 1.2 at left and *v*_*sh*_ = 0.9 at right. (c) Sleep onset and sleep offset timing versus *v*_*sh*_ (left) and *v*_*wc*_ (right).

Together, these results reveal a nonlinear and synergistic interaction between circadian and homeostatic drives, mediated by parallel inhibitory and excitatory circadian pathways. Notably, comparable sleep–wake patterns can emerge from distinct combinations of (*v*_*wc*_, *v*_*sh*_), reflecting parameter degeneracy in the underlying physiology. While such degeneracy confers robustness to the sleep–wake system, it complicates the identification of biologically specific parameter regimes from behavioral observations alone, underscoring the need for integrating additional empirical constraints and data-driven inference in individualized sleep modeling and management.

### Sleep Deprivation as a Probe of Circadian-Modulated Thresholds

Sleep deprivation provides a stringent and physiologically meaningful perturbation of the sleep-wake system, pushing it far from its nominal operating regime and directly testing the stability, resilience, and recovery properties of the underlying dynamics [32, 48]. Within our framework, such extreme conditions offer a unique opportunity to interrogate the functional role of circadian-modulated switching thresholds and to constrain otherwise degenerate parameter regimes. In the threshold formulation developed above, prolonged wakefulness drives the homeostatic variable well above the circadian-modulated sleep-onset threshold, while recovery sleep reflects the subsequent re-crossing of dynamically evolving wake thresholds under circadian control. We therefore treat sleep deprivation not as a parameter-fitting exercise, but as a controlled dynamical excursion through the threshold geometry of the system. Specifically, we examine whether the model reproduces two robust experimental hallmarks of recovery sleep [32]: (i) *Enter Sleep* (ES), characterized by immediate sleep onset following acute, prolonged deprivation (e.g., ∼ 80 h of sustained wakefulness), and (ii) *First Night Recovery* (FNR), marked by partial recovery sleep during the first night after chronic deprivation, compensating approximately 10–30% of the accumulated sleep loss following ∼ 60 h of deprivation.

We first consider the 80 h deprivation paradigm (Fig. 4a). For low values of the circadian wake gain *v*_*wc*_, the transitions immediately into sleep once enforced wakefulness is released (upper panel), reproducing ES phenotype. In contrast, for sufficiently large *v*_*wc*_, sleep initiation fails altogether (lower panel), despite extreme accumulation of homeostatic pressure. This breakdown arises from a circadian elevation of the sleep-onset threshold (circled in Fig. 4b, bottom panel), caused by strong excitatory input to wake-promoting Orx/LC neurons. Even extreme homeostatic load is then insufficient to trigger the wake-to-sleep transition. The dynamical mechanism becomes geometrically transparent in the (*D*_*w*_, *D*_*s*_) plane (Fig. 4c). For small *v*_*wc*_, the post-deprivation trajectory crosses the sleep boundary *D*_*s*_ = *f*_1_(*D*_*w*_) at deprivation offset, triggering an immediate transition into sleep. For large *v*_*wc*_, the trajectory remains confined to the wake domain, never intersecting the sleep threshold despite maximal homeostatic load. Thus, satisfaction of the ES criterion requires *v*_*wc*_ to lie below a critical upper bound, which is also derived in our theoretical analysis (Eq. 17 and dashed line in Fig. 2e).

**Fig 4.**
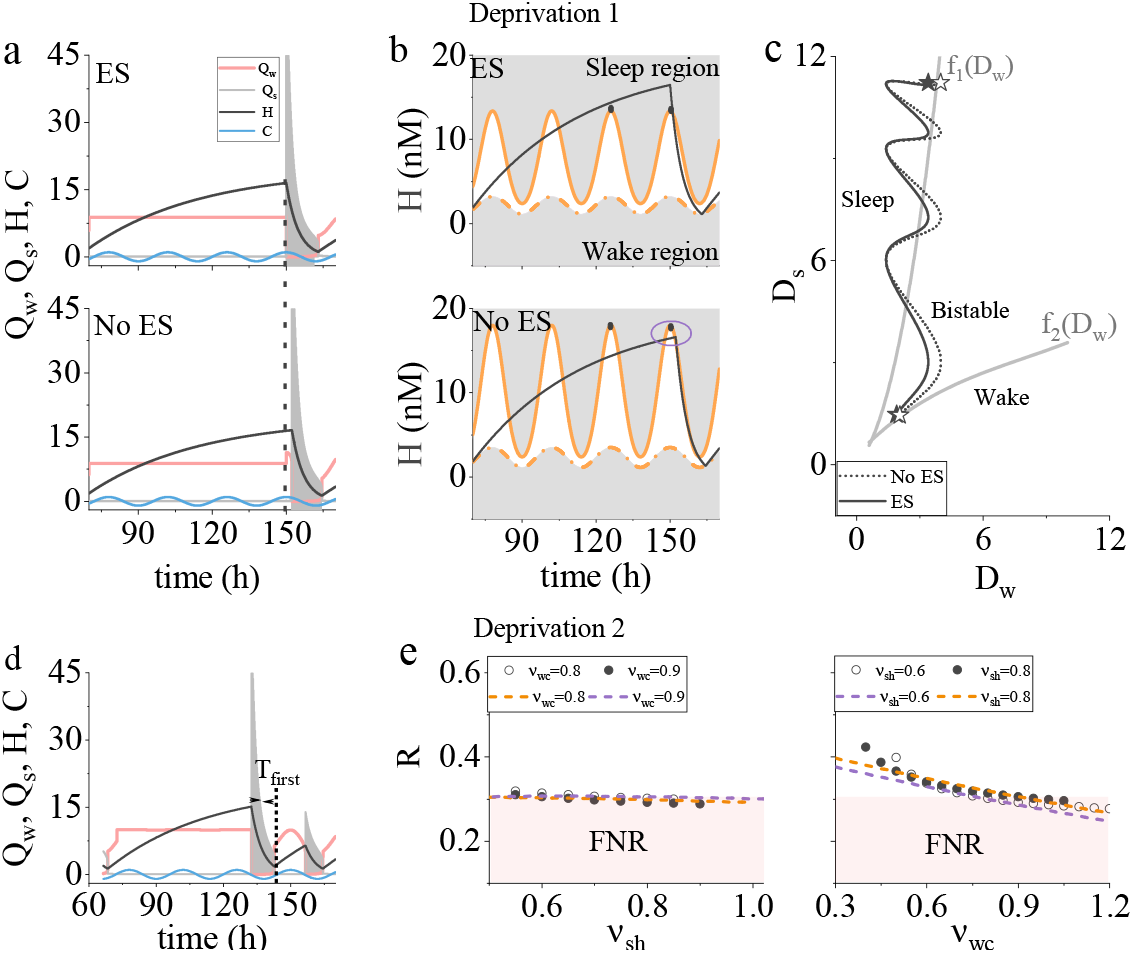
Model behaviors under sleep deprivations. (a) Model response to a 80h sleep deprivation for *v*_*sh*_ = 0.7 with two values of *v*_*wc*_: 1.0 (top) and 1.3 (bottom). Parameters are denoted by hollow and solid stars in Fig. 2e. Dashed line indicates deprivation end, aligned with circadian boundary peak. (b) Time course of homeostatic pressure (black lines), sleep threshold (orange solid lines), and wake threshold (orange dash-dotted lines). “ES” and “No ES” denote the presence or absence of immediate sleep onset following deprivation. (c) Trajectories during the deprivation period in the (*D*_*w*_, *D*_*s*_) plane. Markers indicate the corresponding parameter sets in Fig. 2e. (d) Model output following a 60h deprivation initiated at habitual sleep onset, shows first-night recovery sleep at *v*_*wc*_ = 1.0 and *v*_*sh*_ = 0.7. (e) Recovery sleep duration ratio *R* versus *v*_*sh*_ (left) and *v*_*wc*_ (right). Dashed lines: analytical results given in Eq. 18.

We next consider the 60 h deprivation paradigm (Fig. 4d). The model produces extended rebound sleep, but the fraction of recovered sleep during the first night (FNR, Eq. 20) depends sensitively on the balance between circadian and homeostatic gains (Fig. 4e). Increasing *v*_*wc*_ elevates the wake threshold and promotes earlier termination of sleep, thereby reducing the ratio *R*, while increasing the homeostatic gain *v*_*sh*_ sustains activation of sleep-promoting populations and prolongs recovery sleep, maintaining *R* within or above the experimentally observed 10–30% range. Hence, satisfying the FNR criterion imposes a lower bound on *v*_*wc*_, which is also derived in our theoretical analysis (Eq. 21 and dotted line in Fig. 2e), since insufficient circadian wake drive leads to unrealistically prolonged recovery. Taken together, the ES and FNR conditions confine *v*_*wc*_ to a narrow intermediate range (red-bounded region in Fig. 2e): it must be small enough to permit sleep initiation after extreme deprivation, yet large enough to prevent excessive rebound sleep. These constraints cannot be inferred from nominal sleep-wake cycles alone, but emerge naturally when the system is driven across multiple circadian-modulated thresholds by extreme perturbation.

Overall, these results demonstrate that circadian-modulated sleep and wake thresholds quantitatively govern both the initiation and the extent of recovery sleep following deprivation. By framing sleep loss as a dynamical excursion relative to moving circadian thresholds, the model unifies immediate sleep onset, partial recovery, and parameter identifiability within a single geometric picture. This perspective provides a principled basis for interpreting rebound sleep, vulnerability to circadian misalignment, and individual differences in recovery from sleep deprivation.

### Sleepiness Predictions under Sleep Restriction/Extension

Quantifying sleepiness is essential for translating subjective experience into objective and comparable measures relevant to clinical assessment, fatigue management, and safety-critical settings such as transportation and shift work [49, 50]. Most existing approaches rely on empirical proxies such as prior sleep duration or circadian phase [51, 52]. By contrast, a mechanistic definition is valuable because it exposes the underlying dynamical structure of sleep-wake regulation and enables predictive metrics that are physiologically interpretable and transferable across protocols. In our framework, the explicit geometry of circadian-modulated sleep-wake thresholds naturally yields a state variable for sleepiness. We define sleepiness as the signed distance between the instantaneous homeostatic drive and the active circadian-dependent sleep-onset boundary, *H*− *H*^+^ (Fig. 5a), a geometric concept originally proposed by Daan *et al*. [3]. This quantity measures how close the system is to an imminent wake-to-sleep transition and therefore integrates accumulated sleep pressure and time-of-day–dependent circadian gating into a single dynamical variable.

**Fig 5.**
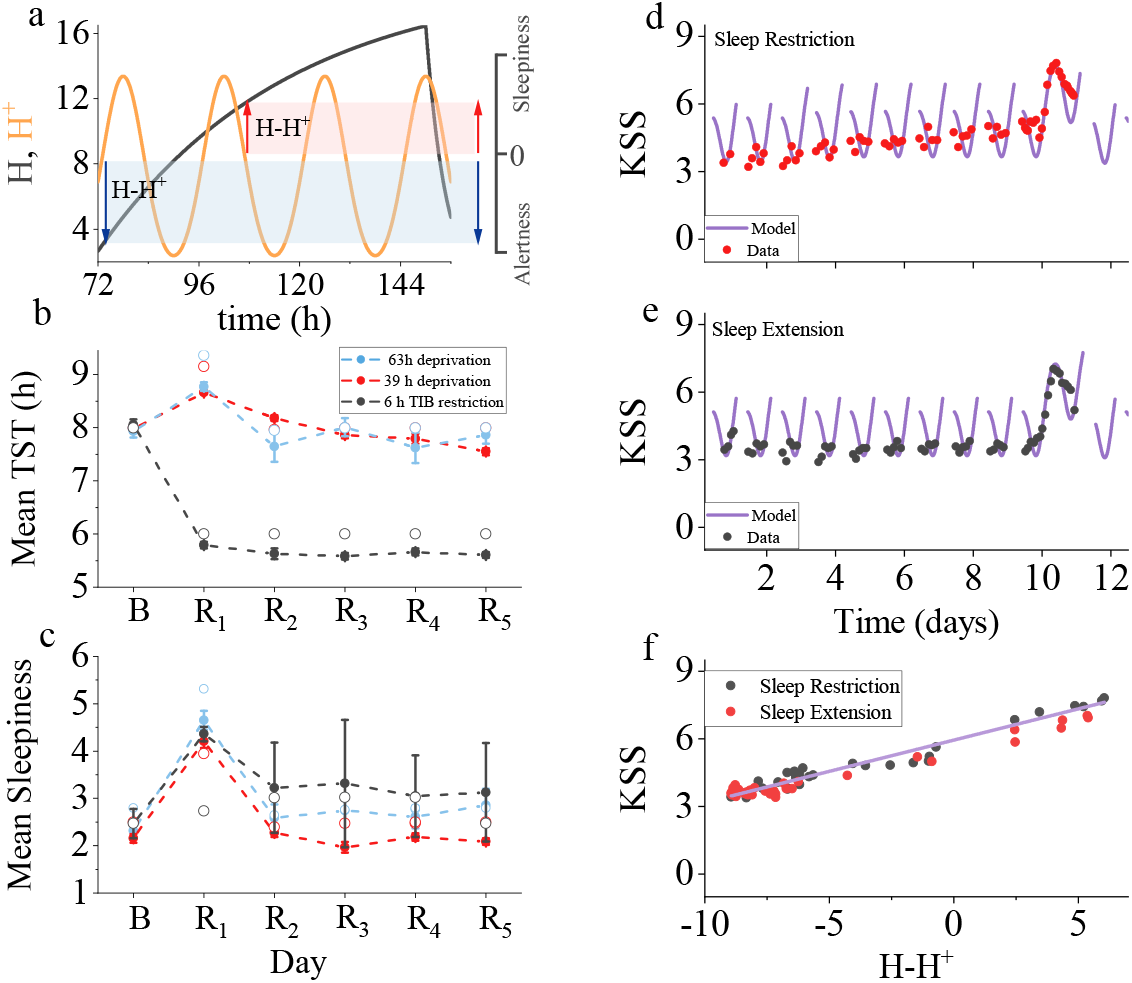
Model fits to behavior data under sleep restriction and extension after sleep deprivation. (a) Schematic illustration of the sleepiness metric ⟨*H*−*H*^+^ ⟩. (b–c) Fits to a dataset comprising three conditions: 39 h sleep deprivation followed by 9 h time in bed (TIB; red), 63 h sleep deprivation followed by 9 h TIB (blue), and 39 h sleep deprivation followed by 6 h TIB (black) [53]. (b) Total sleep time (TST) during recovery; open circles denote model predictions. (c) Mean sleepiness scores for the three protocols; open circles denote model-predicted values obtained via a linear mapping of ⟨*H*− *H*^+^ ⟩. (d–f) Fitting to an independent protocol involving sleep restriction or extension followed by prolonged sleep deprivation [54]. Time courses of KSS scores and the corresponding fitted *H* − *H*^+^ for sleep restriction (d) and extension (e). (f) KSS versus *H* − *H*^+^.

We first evaluated this metric using an experimental protocol comprising multiple deprivation paradigms (63 h or 39 h sleep deprivation followed by 9 h time-in-bed (TIB) extension, and 63 h sleep deprivation followed by 6 h TIB restriction) followed by five recovery nights (R1-R5) with ∼ 9 h TIB [53]. The model accurately reproduced total sleep time (TST) throughout the protocol (Fig. 5b). Crucially, mean KSS scores were well captured by simulated averages of *H*−*H*^+^ (Fig. 5c), providing cross-paradigm validation that the threshold distance tracks subjective sleepiness under both acute and chronic sleep loss. We further tested this relationship using an independent protocol involving systematic sleep restriction or extension followed by prolonged constant-routine deprivation [55, 56]. After a habituation and a baseline night, participants underwnet seven nights of restricted (∼ 6 h TIB) or extended sleep (∼ 10 h TIB), followed by a 39∼ 41 h constant-routine period with repeated KSS assessments [54]. Replicating this design in simulations and sampling *H*− *H*^+^ at the corresponding times revealed a robust linear relationship between the model variable and KSS (Figs. 5d–f):

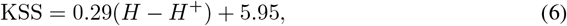

as shown in Fig. 5f. Strikingly, the same mapping held across markedly different sleep-loss histories, including acute deprivation, chronic restriction, sleep extension, and recovery.

This protocol-invariant linear mapping is nontrivial: it implies that subjective sleepiness is not determined directly by sleep history or clock time, but by the instantaneous position of the system relative to its circadian-modulated transition boundary. In other words, different deprivation or restriction schedules converge onto a common low-dimensional physiological state, captured by *H*− *H*^+^, which governs perceived sleepiness. The threshold distance therefore serves not only as a predictive variable, but also as a mechanistic coordinate of the sleep-wake regulation system. This interpretation enables a direct link between subjective sleepiness and its underlying physiological determinants. As the sleep-onset boundary *H*^+^ is an explicit function of circadian drive and neural coupling strengths, and *H* evolves according to simple homeostatic dynamics, the quantity *H*−*H*^+^ provides a transparent decomposition of sleepiness into circadian and homeostatic contributions.

Motivated by this structure, we now examine how model parameters shaping circadian excitation and homeostatic accumulation regulate the average level of sleepiness. To this end, we analyzed the time-averaged index ⟨*H*− *H*^+^ ⟩across parameter regimes (Fig. 6). Increasing the homeostatic gain elevates ⟨*H*− *H*^+^ ⟩, reflecting stronger impact (Fig. 6a) of sleep pressure. Conversely, increasing circadian excitation of wake-active neurons systematically decrease ⟨*H*−*H*^+^ ⟩ (Fig. 6b), consistent with enhanced alerting effects mediated by Orx and LC pathways. Besides, Increasing the accumulation rate of sleep pressure also elevates ⟨*H*− *H*^+^⟩, reflecting stronger accumulation (Fig. 6c) of sleep pressure. Across parameter sweeps, numerical simulations closely follow analytical predictions, supporting the interpretation of sleepiness as a direct consequence of circadian-modulated thresholds. The resulting analytical expression as given in Eq. 26 therefore provides a compact and mechanistically transparent link between subjective sleepiness and its governing physiological parameters, including homeostatic dynamics, circadian coupling strengths, and prior wake duration.

**Fig 6.**
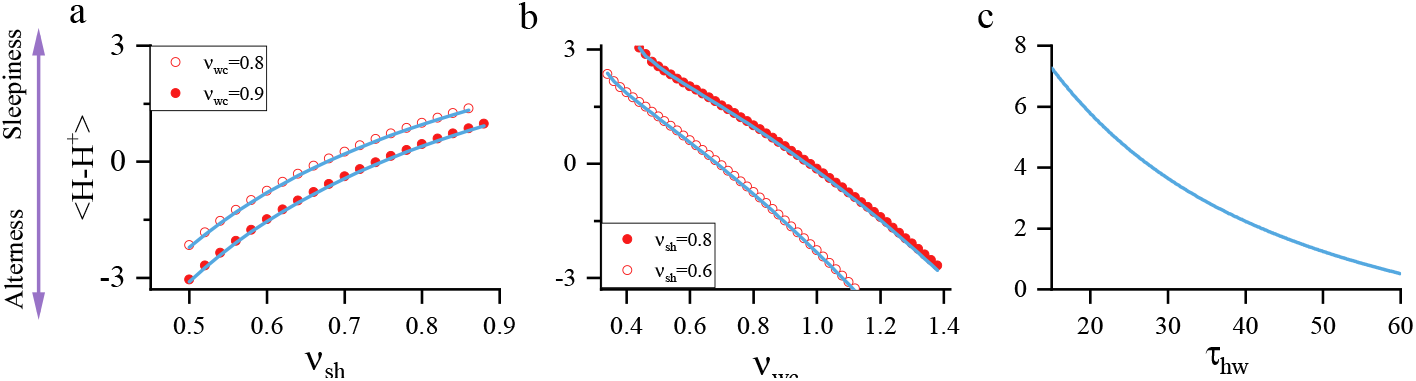
Several factors that affect sleepiness metric ⟨*H*−*H*^+^ ⟩ : *v*_*sh*_, *v*_*wc*_, an d *τ*_*hw*_. Circles and lines in (a-b) denote the simulation and analytical results given in Eq. 26, respectively.

Together, these results establish circadian-modulated threshold distance as a unifying, physiologically grounded metric that links neural population dynamics to subjective sleepiness across protocols. This formulation provides a principled basis for predicting vulnerability to fatigue under sleep restriction, circadian misalignment, and extended wakefulness, and offers a foundation for future integration with objective performance measures in clinical and operational settings.

## Discussions

In this work, we developed a modified PR model that introduces *circadian-dependent sleep and wake thresholds*, providing a mechanistic account of how circadian drive (Process C) and homeostatic pressure (Process S) jointly govern sleep-wake transitions. By constraining the model using normal sleep duration and two independent post-deprivation criteria, we identify a physiologically plausible parameter regime that requires not only sufficient homeostatic sleep pressure but also adequate excitatory circadian input to wake-active populations via the Orx/LC pathway.

Through a combination of bifurcation analysis, numerical simulations, and analytical reductions, we clarify that circadian and homeostatic processes act as separable yet interacting control dimensions governing state stability and switching. Their coupling simultaneously accounts for stable daily sleep-wake cycling, immediate and partial recovery following sleep deprivation, and systematic, protocol-dependent variations in subjective sleepiness. Compared with models based on static thresholds or empirical sleep-propensity curves, our framework links behavioral outcomes directly to identifiable neural pathways, yielding a theory of sleep-wake regulation that is more physiologically grounded, analytically tractable, and operationally predictive.

### Dynamical mechanism of Circadian-Homeostatic coupling for sleep-wake cycles

In the absence of circadian input, increasing homeostatic gain drives the system from sustained wakefulness to monophasic and polyphasic sleep, and ultimately to stable sleep–wake oscillations through a SNIC bifurcation. This establishes that rhythmic alternation can arise intrinsically from homeostatic feedback alone. Introducing circadian input to either sleep-promoting or wake-promoting populations entrains these oscillations to a 24-h rhythm, with classical Arnold-tongue synchronization regions defining the conditions for stable entrainment. Physiologically realistic sleep–wake patterns require sufficient contributions from both processes; weakening either circadian drive or homeostatic accumulation leads to fragmented or unstable dynamics. Robust sleep–wake organization therefore does not arise from circadian forcing alone, but emerges from structured nonlinear interaction between the two processes.

A central contribution of this study is the analytical derivation of circadian-modulated sleep and wake thresholds from the fast subsystem. These thresholds formalize when transitions occur and expose how circadian and homeostatic signals jointly shape state stability. Stronger circadian excitation of wake-active populations elevates the sleep-onset threshold, delays both sleep onset and offset, consolidates wakefulness, and shortens total sleep duration. At the same time, multiple combinations of circadian and homeostatic gains can produce similar macroscopic sleep patterns, revealing substantial parameter degeneracy. This finding underscores the importance of physiological and behavioral constraints–such as deprivation responses and sleepiness dynamics–for identifying biologically plausible regimes.

### Sleep deprivation, recovery, and sleepiness as dynamical probes

Sleep deprivation provides a stringent test of sleep–wake models because it pushes the system far from its nominal operating regime. Within the threshold framework, deprivation corresponds to a large excursion of the homeostatic variable relative to circadian-modulated transition boundaries. The model reproduces two robust experimental hallmarks: immediate sleep onset after extreme deprivation and partial first-night recovery following chronic deprivation. Elevated circadian excitation delays post-deprivation sleep onset and limits rebound, whereas weaker circadian excitation or stronger homeostatic coupling prolongs recovery sleep. These results demonstrate that recovery behavior is governed by the same threshold geometry that controls normal state switching and provides quantitative constraints on circadian excitation of wake-promoting circuits. Deprivation and recovery therefore emerge naturally from the same dynamical structure, rather than requiring hoc mechanisms.

We further quantified sleepiness as the distance between homeostatic drive and the active circadian sleep boundary, turning subjective alertness into an objective, model-based state variable. Across diverse laboratory protocols–including sleep restriction, sleep extension, acute deprivation, and recovery–this variable mapped linearly onto KSS scores. Importantly, this mapping held independently of how sleep debt was accumulated, indicating that subjective sleepiness reflects the instantaneous dynamical state of the circadian-homeostatic system rather than sleep history per se. As the sleep boundary can be given an explicit function of circadian phase and neural coupling strengths, this metric directly links subjective experience to physiological control parameters. Analytical approximations further relate average sleepiness to homeostatic time constants, coupling gains, circadian drive strength, and deprivation duration, providing a transparent bridge between phenomenology and mechanism. This offers a principled alternative to empirical fatigue models based solely on prior sleep or clock time.

### Clinical, operational, and modeling implications

Introducing dynamic, circadian-modulated sleep and wake thresholds represents a substantial advance over traditional models based on static or heuristic transition criteria. By embedding circadian modulation directly into the switching mechanism, the framework enables circadian misalignment and its behavioral consequences to be quantified mechanistically, while treating endogenous rhythms, environmental perturbations, and neural circuitry within a single dynamical system.

This perspective provides a unified interpretation of several well-known phenomena. The wake maintenance zone (WMZ) [12, 57], for instance, emerges naturally when circadian excitation of Orx/LC neurons transiently elevates the sleep-onset threshold, stabilizing evening wakefulness despite high homeostatic pressure. The model predicts that the depth and timing of this wake fate covary with circadian amplitude and Orx/LC integrity, offering a mechanistic explanation for inter-individual variability and for its compression in aging or neurological conditions. In older adults, reduced circadian amplitude and altered arousal circuitry weaken circadian stabilization of wakefulness, leading to early sleep onset, fragmented sleep, and increased evening sleep intrusions [58]. Within the threshold framework, these features arise from the same altered transition geometry that governs normal sleep-wake switching, without invoking separate age-specific mechanisms.

For shift workers, circadian misalignment displaces sleep and wake boundaries into biologically unfavorable phases, producing delayed sleep onset, early awakening, curtailed sleep duration, and pronounced night-shift sleepiness that cannot be predicted from time awake alone [59]. Circadian-modulated thresholds naturally capture these effects and suggest that interventions targeting sleep timing, light exposure, and circadian phase may be more effective than strategies focused solely on sleep duration [20]. More broadly, our results indicate that misalignment between circadian and homeostatic processes–whether due to shift work, jet lag, aging, or genetic variation–can be understood as a distortion of transition geometry in state space. Restoring appropriate alignment of circadian-modulated thresholds with homeostatic dynamics therefore constitutes a principled target for improving sleep health and performance.

Operationally, the approximately linear mapping between threshold distance and clinical sleepiness scores enables translation from a mechanistic state variable to a widely used subjective measure. This allows sleepiness to be predicted under untested schedules–such as rotating shifts, split sleep, or irregular duty cycles–without protocol-specific calibration [60]. Extending the framework to objective vigilance metrics, such as Psychomotor Vigilance Test (PVT) [61], would further unify prediction of perceived and functional impairment in safety-critical settings, including transportation and industrial operations.

### Limitations and future directions

Several limitations point to directions for future work. First, environmental and lifestyle factors such as light exposure, diet, and exercise are not explicitly modeled; incorporating these influences could improve ecological validity and support personalized, non-pharmacological interventions. Second, individual variability in circadian dynamics, arousal circuitry, and chronotype is not yet represented and will be important for population-level and clinical applications. Third, further experimental validation using physiological recordings and behavioral data will be essential to constrain parameter ranges and test model predictions in real-world settings.

## Methods

### Analytical derivation of sleep-wake transition thresholds

Here we derive sleep-wake transition thresholds directly from the steady states of the wake-active and sleep-active neural populations in the fast subsystem of the model. At equilibrium, the mean membrane potentials of the wake 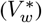 and sleep 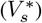 populations satisfy two coupled self-consistency equations determined by recurrent inhibition, sigmoid firing responses, and external drives to each population:

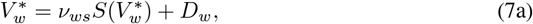

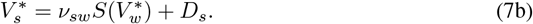

Solving these equations is equivalent to finding the zeros of an implicit scalar function

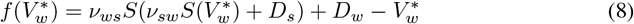

and an analogous expression for 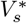. Transitions between behavioral states occur when stable and unstable equilibria collide through saddle-node bifurcations. Accordingly, critical transition boundaries are obtained by solving simultaneously for 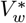at which the implicit function vanishes 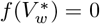 and its derivative 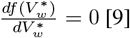.

This procedure yields two analytical threshold curves that relate the effective sleep drive *D*_*s*_ to the wake drive *D*_*w*_:

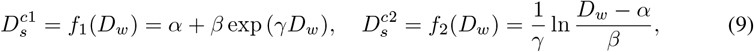

with coefficients *α* = −3.84, *β* = 3.38, *γ* = 0.39. The first curve, *f*_1_, defines the boundary for sleep onset, and the second curve, *f*_2_, defines the boundary for awakening. Both curves take closed exponential-logarithmic forms, parameterized by three fitted constants obtained from the neural activation function.

Substituting the explicit circadian and homeostatic contributions into *D*_*w*_ and *D*_*s*_ produces explicit, circadian-modulated sleep pressure thresholds:

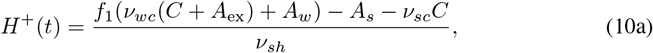

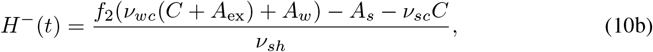

representing the amount of homeostatic sleep pressure required to trigger sleep onset (*H*^+^) and awakening (*H*^*−*^) at given circadian phase. In the model, sleep occurs when *H*(*t*) exceeds *H*^+^(*t*), and wake occurs when *H*(*t*) falls below *H*^*−*^(*t*).

Under periodic circadian forcing, the system settles into a 24-h switching cycle. Within each cycle, homeostatic pressure accumulates during wake until it reaches *H*^+^, triggering sleep, and decays during sleep until it reaches *H*^*−*^, triggering awakening. Enforcing periodicity yields a closed set of relations linking sleep onset time, sleep duration, and the two thresholds:

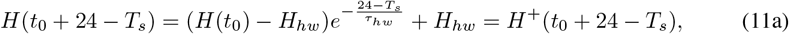

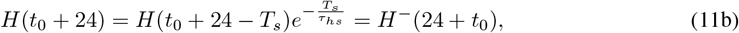

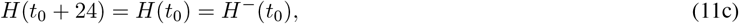

where 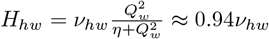 denotes the saturation level of homeostatic pressure during sustained wakefulness, *t*_0_ denotes the sleep offset time and *T*_*s*_ is sleep duration. Solving these equations analytically provides an explicit approximation for sleep duration and circadian phase at sleep onset and offset as a function of physiological parameters, which were validated in Figs. 3(b–c).

*Derivation details*. From the periodicity conditions of the sleep-wake cycle, the awakening threshold at time *t*_0_ satisfies

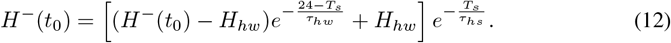

For physiological plausible sleep durations (6 ∼10 h), the exponential accumulation factor during wake, 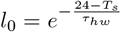, varies only weakly (approximately 0.6703 ∼ 0.7326) and can therefore be well approximated by a constant, *l*_0_ ≈ 0.7. This approximation yields a compact analytical of sleep duration

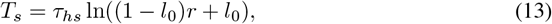

where the ratio 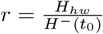, and *H*^*−*^(*t*_*0*_) is given in Eq. 10b. It makes explicit how sleep duration is jointly determined by the underlying physiological parameters.

### Analytical responses to sleep deprivations

As the thresholds depend explicitly on circadian phase and homeostatic pressure, responses to enforced wakefulness can be solved analytically and compared directly with numerical simulations (Fig. 2e and Fig. 4e).

### ES Criterion: immediate sleep following prolonged deprivation

To model enforced wakefulness, we prevent state switching and maintain wake dominance *Q*_*w*_ *> Q*_*s*_ for a duration *T*_*sd*_ (*T*_*sd*_ ≥ 40 h). During this interval, homeostatic pressure increase exponentially toward its saturation level as

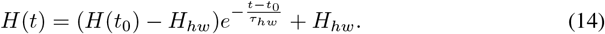

At deprivation termination, sleep onset is required to occur even when circadian wake drive is maximal (*C* = 1) [32]. This condition imposes that the accumulated homeostatic pressure exceed the circadian-elevated sleep threshold evaluated at peak circadian phase

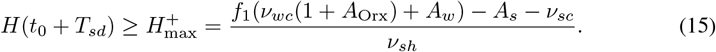

Rearranging this condition yields: 1) an explicit lower bound on the homeostatic gain *v*_*sh*_:

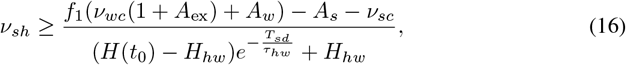

ensuring that sleep pressure can overcome circadian arousal; 2) an upper bound on circadian excitation of wake-active neurons *v*_*wc*_:

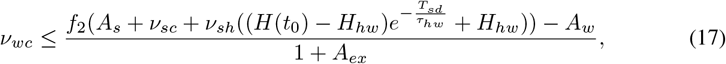

preventing excessive stabilization of wakefulness. These inequalities explain analytically why overly strong circadian excitation of the Orx/LC pathway can block sleep even under extreme sleep debt. Our analytical predictions (Fig. 2e, dashed line) closely match numerical simulations, where parameter values *t*_0_ = 70 h and *T*_*sd*_ = 80 h were chosen so that deprivation ended near circadian peak, i.e., *C*(*t*_0_ + *T*_*sd*_) = 1, providing the most stringent condition for sleep initiation with *H*(*t*_0_) ≈ 2.

### FNR Criterion: first-night recovery sleep duration

For deprivation beginning at the normal sleep offset *t*_0_ with a duration *T*_*sd*_ ≥ 40 h, homeostatic pressure at recovery onset *H*_1_ = *H*(*t*_0_ + *T*_*sd*_) can be expressed analytically as a weighted sum of its initial value and its saturation level. Recovery sleep terminates when the decaying homeostatic pressure *H*(*t*_0_ + *T*_*sd*_ + *T*_first_) reaches the circadian-dependent awakening threshold *H*^*−*^. Solving this condition yields an explicit expression for first-night recovery sleep duration as a function of homeostatic parameters, circadian phase at recovery end, and circadian and homeostatic coupling strengths, as

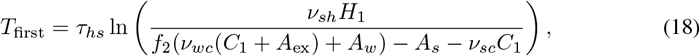

where *C*_1_ = sin(*ω*_*c*_(*t*_0_ + *T*_*sd*_ + *T*_first_)). Numerical results (Fig. 4e) validate this analytical prediction.

This formula predicts a physiologically plausible rebound range in which recovery sleep exceeds baseline sleep by approximately 10 ∼ 30% to be consistent with empirical observations [32], given as

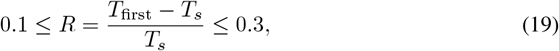

where *T*_*s*_ and *T*_first_ are given in Eq. (18) and Eq. (13), respectively. So we can obtain

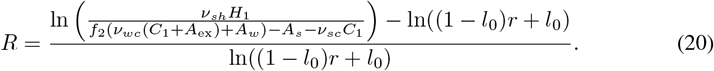

Imposing the bound *R*≤ 0.3 and using the formula of normal and recovery sleep duration, Eq. 13 and Eq. 18, yields a complementary constraint: a lower bound on circadian excitation strength, given as

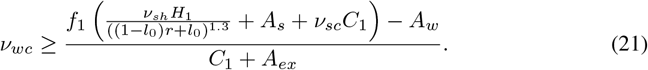

Our analytical predictions (Fig. 2e, dotted line) closely match numerical simulations across deprivation protocols, with *T*_*sd*_ = 60 h, *C*_0_ = −0.865 and *C*_1_ = −0.3. Together with the ES criterion, these conditions define a narrow physiologically plausible parameter regime.

### Sleepiness prediction

Sleepiness (KSS metric) is quantified as the time-averaged distance between homeostatic sleep pressure and the circadian-modulated sleep-onset threshold:

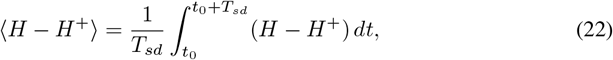

where the threshold is given by

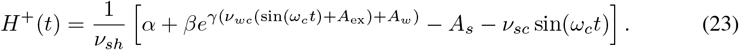

This quantity links subjective sleepiness directly to the system’s dynamical distance from the bifurcation-defined boundary for sleep onset *H*^+^(*t*). It provides a mechanistic measure of sleepiness: the distance decreases when homeostatic pressure accumulates toward the threshold or when circadian excitation elevates the sleep threshold.

As *H*^+^(*t*) explicitly incorporates circadian excitation and inhibition through *v*_*wc*_ and *v*_*sc*_, while *H*(*t*) evolves on the slow homeostatic timescale, ⟨*H*− *H*^+^ ⟩captures the joint effect of circadian and homeostatic processes on subjective sleepiness. To obtain explicit parameter dependence, we approximate ⟨*H*− *H*^+^ ⟩using a Fourier–Bessel expansion of the circadian exponential term *e*^*a* sin (*ϕ*)^. This yields an closed-form expression (Eq. 26) that links mean sleepiness to: 1) homeostatic parameters *τ*_*hw*_, *v*_*sh*_ and *H*_*hw*_; 2) circadian coupling strengths *v*_*wc*_ and *v*_*sc*_; and 3) sleep deprivation duration *T*_*sd*_.

This result provides a direct mechanistic bridge between physiology, nonlinear dynamics, and subjective experience: by expressing sleepiness in terms of circadian-modulated transition thresholds, the same framework that accounts for sleep-wake transitions and recovery after deprivation also predicts alertness regulation. Thus, sleepiness becomes a state variable of a single coherent dynamical system, enabling principled assessment of vulnerability to fatigue under sustained challenge.

*Derivation outline*. As *H*(*t*) lacks a closed-form anti-derivative, we expand *e*^*a* sin (*ϕ*)^ as:

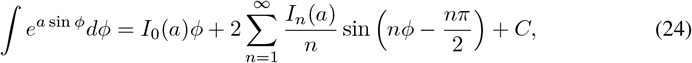

with 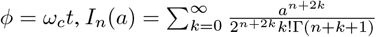, where Γ is the Gamma function. We further approximate *I*_*n*_(*a*) as

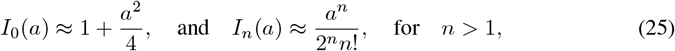

leading to

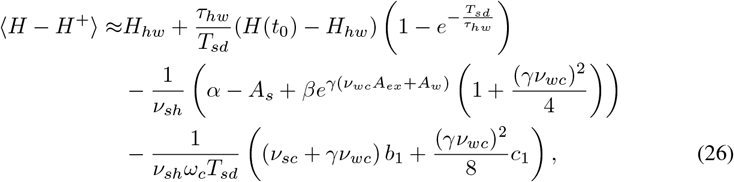

With

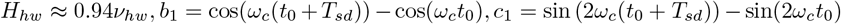

Equation 26 shows that the mean deviation between the instantaneous homeostatic drive and its up-threshold value, ⟨*H*−*H*^+^ ⟩, can be decomposed into three interpretable components: (i) a purely homeostatic relaxation term, reflecting exponential convergence toward the intrinsic lower bound *H*_*hw*_ over the window *T*_*sd*_; (ii) a tonic shift determined by the slow–wake contribution from *α*− *A*_*s*_ and the exponentially weighted excitation gain 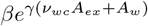, which together scale the long-term displacement of *H* from *H*^+^; and (iii) an oscillatory correction proportional to 1*/ω*_*c*_*T*_*sd*_, which captures the net effect of periodic modulation on short timescales. The small sinusoidal terms, governed by *b*_1_ and *c*_1_, decay rapidly with increasing frequency or averaging window, confirming that rhythmic perturbations only weakly bias the cycle-averaged homeostatic level. Notably, the leading Bessel contribution tightens this analytical estimate, enabling accurate tracking of mean dynamics even when the wake input contains sizable rhythmic structure.

Collectively, these results demonstrate that the model predicts a well-behaved,frequency-robust average homeostatic load, with oscillatory corrections remaining subdominant across physiologically relevant parameter ranges. This analytical predictions are well confirmed in numerical simulations (Fig. 6).

## Acknowledgments

This work was supported partially by the Natural Science Foundation of Zhejiang Province under Grant Nos. LY24A050003, Jiaxing Public Welfare Project under No. 2025CGZ037, the Humanities and Social Science Fund of Ministry of Education of China under Grant Nos.25YJAZH212, the National Natural Science Foundation of China under Grant Nos. 12175242, and Ministry of Science and Technology of the People’s Republic of China (STI2030-Major Projects2021ZD0201900), National Natural Science Foundation of China (grant Nos. 12090052 and U24A2014).

